# Experimental Evolution *in vivo* to Identify Selective Pressures During Pneumococcal Colonization

**DOI:** 10.1101/2020.01.15.908590

**Authors:** Vaughn S. Cooper, Erin Honsa, Hannah Rowe, Christopher Deitrick, Amy R. Iverson, Jonathan J. Whittall, Stephanie L. Neville, Christopher A. McDevitt, Colin Kietzman, Jason W. Rosch

## Abstract

Experimental evolution is a powerful technique to understand how populations evolve from selective pressures imparted by the surrounding environment. With the advancement of whole-population genomic sequencing it is possible to identify and track multiple contending genotypes associated with adaptations to specific selective pressures. This approach has been used repeatedly with model species *in vitro*, but only rarely *in vivo*. Herein we report results of replicate experimentally evolved populations of *Streptococcus pneumoniae* propagated by repeated murine nasal colonization with the aim of identifying gene products under strong selection as well as the population-genetic dynamics of infection cycles. Frameshift mutations in one gene, *dltB*, responsible for incorporation of D-alanine into teichoic acids on the bacterial surface, evolved repeatedly and swept to high frequency. Targeted deletions of *dltB* produced a fitness advantage during initial nasal colonization coupled with a corresponding fitness disadvantage in the lungs during pulmonary infection. The underlying mechanism behind the fitness tradeoff between these two niches was found to be enhanced adherence to respiratory cells balanced by increased sensitivity to host-derived antimicrobial peptides, a finding recapitulated in the murine model. Additional mutations were also selected that are predicted to affect trace metal transport, central metabolism and regulation of biofilm production and competence. These data indicate that experimental evolution can be applied to murine models of pathogenesis to gain insight into organism-specific tissue tropisms.

**Importance:** Evolution is a powerful force that can be experimentally harnessed to gain insight into how populations evolve in response to selective pressures. Herein we tested the applicability of experimental evolutionary approaches to gain insight into how the major human pathogen *Streptococcus pneumoniae* responds to repeated colonization events using a murine model. These studies revealed the population dynamics of repeated colonization events and demonstrated that *in vivo* experimental evolution resulted in highly reproducible trajectories that reflect the environmental niche encountered during nasal colonization. Mutations impacting the surface charge of the bacteria were repeatedly selected during colonization and provided a fitness benefit in this niche that was counterbalanced by a corresponding fitness defect during lung infection. These data indicate that experimental evolution can be applied to models of pathogenesis to gain insight into organism-specific tissue tropisms.

## Introduction

*Streptococcus pneumoniae* (the pneumococcus) typically exists as a commensal member of the nasal flora, colonizing individuals for extended periods of time without the development of invasive disease. Competition within the nasopharynx between bacterial species is complex, with both synergistic and antagonistic relationships between bacterial species^1–7^. Critical to the success of the pneumococcus as a human pathogen is its capacity to transmit to a new host and establish colonization. This is one of the major selective pressures on the pneumococcus, as invasive disease is typically envisioned as an evolutionary dead-end due to clearance by the immune system or antibiotics or via the death of the host, thereby preventing further transmission of the strain. As such, the pneumococcus has evolved numerous strategies to facilitate efficient transmission and colonization of new hosts^8–13^. Mutational screens have identified many pneumococcal genes that are required for successful colonization of the nasal passages as well as for invasive infection of the lungs and the bloodstream^14,15^. One potential limitation of transposon screens is they produce loss of function mutations that mostly identify genetic deficits, whereas naturally arising genetic variation is dominated by SNPs that can produce a range of phenotypes^16,17^. Colonization of any new environment, such as the upper airway, likely selects for new or altered phenotypes. Accordingly, experimental evolution is an ideal method to investigate organism and population-wide adaptation to this niche.

Experimental evolution has proven insightful for many problems in microbiology and evolutionary biology. Large microbial populations will naturally generate ample genetic variation from spontaneous mutation and empower natural selection to enrich beneficial mutations. With replicate bacterial populations propagated under similar conditions, phenotypes and genotypes that recurrently evolve can be identified, and this parallelism is a hallmark of adaptation^18^. This approach has proven invaluable to understand how antibiotic resistance evolves, as well as the factors that drive bacterial transition from planktonic to biofilm states^19–21^. Extending these approaches to bacterial systems has not been extensively explored *in vivo*, but the same evolutionary principles of niche adaptation apply when bacterial populations remain large enough to empower selection on naturally arising mutations that improve niche fitness. Evidence of strong selection in humans comes from immunological screening of IgG binding affinities, which has revealed diverse evolutionary patterns for many of the surface exposed antigens encoded by the pneumococcus^22,23^. These studies indicate the surface properties of the pneumococcus are under strong selective pressure, both in term of functionality and evasion of host immune recognition.

One of the longest studied and immunogenic surface structures of the pneumococcus is the polysaccharide capsule. Introduction of the pneumococcal conjugate vaccine dramatically altered the fitness landscape of the circulating strains whose serotypes were the basis of the vaccine. Capsular serotype is a critical factor in mediating carriage duration and as such the locus encoding the biosynthetic machinery for capsule production displays heightened recombination and substitution frequencies, which can lead to novel serotypes^24,25^. While the capsule is critically important in the evasion of innate immune clearance, the biochemical properties of the polysaccharide capsule also play an important role in pneumococcal colonization and fitness. This has been demonstrated both at a population level and in murine models, where strains encoding more negatively charged capsules, resulting in more negative surface charge potentials, have a competitive advantage during colonization, especially during competition with other pneumococci within the nasal passages^26^. Capsule switching can incur both fitness advantages and disadvantages during colonization and transmission^9,27^. Capsule biosynthesis is linked to a number of additional cellular processes, including metabolism ^28,29^. Transcriptional control of capsule biosynthesis is a critical mediator of invasive disease, a finding also reflected by the polymorphisms observed in the promoter region of the capsule locus^30,31^. These findings underscore the vitally important role of the pneumococcal surface polysaccharide capsule for both immune evasion as well as nasal colonization and invasive disease capacity.

We hypothesized that repeated inoculation and colonization of the nasal passages would select for distinct genotypes of *S. pneumoniae* with enhanced colonization properties. Using whole-population, whole-genomic sequencing and new methods to infer genotype structure, we were able to track individual mutations and the rise of different haplotypes in the pneumococcal population during repeated passages in murine hosts. We identified several parallel, null mutations in *dltB*, responsible for incorporation of D-alanine into teichoic acids on the bacterial surface that was under strong positive selective pressure^10^. Subsequent deletion of *dltB* resulted in a fitness benefit during experimental murine nasal colonization and enhanced bacterial adherence to cultured epithelial cells. This fitness was counterbalanced by an increased sensitivity to host derived antimicrobial peptides and attenuation during experimental pneumococcal pneumonia. These findings indicate that modulation of the surface charge of the pneumococcus is a critical aspect of colonization efficiency and that *in vivo* experimental evolution can be leveraged to understand niche adaptation in bacterial pathogens.

## Materials and Methods

### Media and growth conditions

*S. pneumoniae* strain BHN97 or the xenogeny derivative, BHN97x, was grown on tryptic soy agar (EMD Chemicals, New Jersey) supplemented with 3% sheep blood or in C+Y, a defined semi-synthetic casein liquid media^32^ supplemented with 0.5% yeast extract. Cultures of *S. pneumoniae* were inoculated from frozen stock and incubated at 37 °C in 5% CO_2_.

### *In vivo* Experimental Evolution

*S. pneumoniae* strain BHN97x was grown to an optical density at 620 nm (OD_620_) of 0.4, corresponding to 10^8^ cells/mL, in C+Y and used to infect three independent lineages of mice. Female BALB/cJ mice (Jackson Laboratory, Bar Harbor, ME) aged seven weeks were maintained in BSL2 facilities. All experiments were done under inhaled isoflurane (2.5%). Mice were challenged intranasally with 5 × 10^5^ colony forming units (CFUs) of BHN97x in 100 mL phosphate-buffered saline (PBS) as described previously^33^. The BHN97x challenge strains had been engineered to express luciferase as described^34^. At 72 h post challenge mice were euthanized and bacteria recovered from the nasal passages via retrotracheal lavage with 3 mLs cold PBS. Nasal lavage was centrifuged for 5 min at 300 × g to pellet host cells. Supernatant was transferred and bacterial cells harvested by 15,000xg centrifugation for 5 min. Bacterial pellets were resuspended in 520 μL ThyB and harvested bacterial were plated on triplicate TSA blood agar plates for enumeration of recovery and to expand the population for the subsequent round of infection. Bacterial populations were recovered following overnight incubation at 37 °C in 5% CO_2_ and resuspended in C+Y media. The bacterial suspension was adjusted to OD_620_ = 0.4 to normalize the following round of infections. Bacterial inoculum and total recovered bacterial populations were enumerated for each infection and subsequent recovery by serial dilution and plating. A total of three experimental evolutionary lineages were maintained for a total of ten generations.

### *In vitro* Competitive Index

For measuring in vitro competitive indexes, all strains were mixed in a 1:1 ratio from titered glycerol stocks with 10^6^ CFUs of each strain for the initial passage. Strains were cultivated in CY media with 0.5% glucose as the carbon source at 37 °C in 5% CO_2_ for 6 h. Following outgrowth, strains were serially diluted and differentiated using respective antibiotic plates, supplemented with kanamycin at 400 μg/mL for BHN97x and erythromycin for the respective deletion mutants. Simultaneously, glycerol stocks were made for the second round of passaging. Bacterial enumeration following overnight incubation at 37 °C in 5% CO_2_ was used to determine competitive index measurements. Five independent cultures were maintained for the respective strain combinations.

### Primers/mutant construction

Mutants in TIGR4 were made by PCR-based overlap extension. Briefly, flanking regions 5' and 3' to the target gene were amplified by PCR and spliced to an antibiotic cassette. The final PCR product was transformed into the pneumococcus by conventional methods, replacing the targeted gene with the erythromycin antibiotic cassette. To confirm transformation, primers outside of the transformed region were used for PCR and the region was subsequently sequenced.

### Sequencing

To improve genotyping of evolved populations, the genome of the founding isolate was sequenced on a PacBio RS2 using 5- to 10-Kb fragment libraries loaded on one SMRTcell (Johns Hopkins Sequencing Center). The closed genome sequence is available through NCBI (PRJNA517171). For experimentally evolved population samples, sequencing libraries were prepared from each passage from each lineage, barcoded by using the Nextera kit (Illumina), and pooled in one lane of an Illumina HiSeq2500 for sequencing, as described previously^35^. All evolved isolates are available through NCBI (PRJNA600000).

### Variant calling

The closed genome was used as a reference for the mapping of all subsequent Illumina short reads at a coverage of 300× or more. Reference mapping and the detection of SNPs, indels, and structural variants was conducted by using breseq (v. 0.28) software as described^36 34^. Putative mutations were extracted from raw output and filtered to remove dubious calls on the basis of the following criteria: i) 2% frequency or less, ii) multiple mutations in one locus found exclusively in 1-2 samples, which is evidence of mis-mapped reads to a different organism in the mouse microbiome, iii) erroneous mis-mapping to transposases or known repetitive elements iv) inconsistent calls in low-frequency homopolymers, indels, and tRNAs.

### Genotype inferences from populations and Muller plots

Mutation filtering, allele frequencies, and plotting were done in R v3.5.3. Muller plots were generated using the lolipop package (https://github.com/cdeitrick/lolipop) v0.6 using default parameters. To summarize, these tools predict genotypes and lineages based on shared trajectories of mutations over time and test their probability of nonrandom genetic linkage. Successive evolution of genotypes, or nested linkage, is identified by a hierarchical clustering method. The method also includes customizable filters that eliminate singletons that do not comprise prevalent genotypes. Muller plots were manually color coded by the presence of putative driver mutations within each genotype. Additional mutations that occurred on the background of putative driver mutations can be viewed in the allele frequency plots also generated by this package.

### LL-37 Sensitivity Assays

To measure sensitivity of the strains to the antimicrobial peptide LL-37, a 1:100 back dilution of an OD_620_ culture was added into fresh C+Y supplemented with serial dilutions of LL-37 as previously described^37^. Bacterial outgrowth was measured by measuring OD_620_ at 37 °C/ 5% CO_2_ in a Biotek plate reader for up to 24 h. Assays were run in quadruplicate from two experimental replicates and the data pooled from all replicates.

### Adherence assay

A549 lung epithelial and Detroit nasopharyngeal cells were grown in 6 well plates at 37C in 5% CO_2_ to >80% confluence. Pneumococcal cultures were grown to OD_620_=0.4, washed and then added to eukaryotic cells at 2×10^6^ cfu/well. Three wells were used for each mutant and the assays were repeated at least three times. For adherence assays, cells were incubated 60 minutes with bacteria, a time chosen to minimize internalization of adherent bacteria. After washing three times in PBS, the cells were released from the plate with trypsin, lysed, serial diluted, and subsequently plated on TSA blood agar plates. Colonies grown overnight were counted as bacteria adherent to cells. Experiments were performed in triplicate.

### Ethics Statement

All experiments involving animals were performed with prior approval of and in accordance with guidelines of the St. Jude Institutional Animal Care and Use Committee. The St Jude laboratory animal facilities have been fully accredited by the American Association for Accreditation of Laboratory Animal Care. Laboratory animals are maintained in accordance with the applicable portions of the Animal Welfare Act and the guidelines prescribed in the DHHS publication, Guide for the Care and Use of Laboratory Animals.

### Zeta Potential Measurements

*S. pneumoniae* strains, wild-type and mutant derivatives thereof, were grown in ThyB to mid-log phase, defined as OD_600_ = 0.3. The cells were harvested by centrifugation and washed in PBS three times. Cells were resuspended to a final OD_600_ = 0.3 in PBS and zeta potential measured using Zetasizer Nano ZS 90 (Malvern, UK), equipped with a Helium–Neon laser (633 nm) as a source of light, with the detection at 90 degree scattering angle at room temperature (28 °C). Zeta potential measurements were carried out under identical experimental conditions and all experiments were performed in biological triplicate.

### Capsule shedding assay and capsule blotting

Capsule shedding and blotting were undertaken as previously described^37^. 1 mL of logarithmic culture of pneumococci grown in C+Y or indicated medium was harvested at OD_620_ 0.4 by centrifugation. The pellet was then washed with 1 mL buffer SMM (0.5 M Sucrose, 0.02M MgCl_2_, 0.02 M 2-(N-Morpholino)-ethanesulfonic acid (MES) pH 6.5) and resuspended in 1mL SMM. Either LL-37 (at 4 mg/ml for standard assays, or the indicated concentration) or antibiotics at the indicated concentration were then added and the samples were incubated at 37 °C for 30 min (or indicated time). Samples were then centrifuged 5 min at >14,000 × g, and the supernatant removed and kept for analysis (supernatant fraction). The cell pellet was then resuspended in 975 mL PBS to which 25 mL of 10% (w/v) deoxycholate was added (for LytA deficient strains sufficient recombinant purified LytA was added to allow for bile salt stimulated lysis) and the cells were allowed to lyse for 30 min at 37 °C (Pellet fraction).To remove significant proteinaceous cross reactive species detected by anti-capsule antisera, before analysis both supernatant and pellet fractions were treated with 5 mL proteinase K (Sigma) for 30 min at 37 °C. Samples were boiled in SDS-PAGE sample buffer and 5 mL was electrophoresed on a 0.8% agarose gel in standard 1X Tris-Acetate running buffer and 3 mL of a 20 mg/mL solution of purified capsular polysaccharides was included as a standard in all gels. After electrophoresis samples were transferred from the agarose gel to mixed nitrocellulose ester membranes (HATF, Millipore) via 20x SSC capillary transfer overnight. After transfer membranes were rinsed in 6x SSC, allowed to air dry, and UV crosslinked at 150,000 mJ using a Stratagene UV crosslinker. The membrane was then blocked with PBS containing 0.1% Tween 20 and 5% non-fat milk for 1 h at room temperature and probed with commercially available serotype specific anti-capsular antiserum (Statens Serum Institute (serotype 19F cat 16911) at a 1:15,000 dilution. After washing the membrane was then visualized using secondary HRP conjugated antibodies (Bio-Rad cat 170-6515) (1:30,000) and imaged on a ChemiDoc MP imager (Bio-Rad). For quantitation of relative capsule amounts blot images were analyzed by densitometry using Image Lab software (Bio-Rad).

### Statistical Analysis

Levels of adherence and colonization were compared by Mann-Whitney U test. A *P* <0.05 was considered significant for all experiments. Standard parametric statistics were conducted in GraphPad Prism, and population-genetic statistics were conducted in R v3.5.3.

## Results

### Experimental evolution of enhanced pneumococcal nasal colonization

To gain insight into the traits under selection during pneumococcal colonization, we designed a sequential nasal passaging model in mice. At the outset, three different mice were infected intranasally with *S. pneumoniae* strain BHN97x, which was allowed to colonize for three days and initiate lineages M1-M3. This is a serotype 19F strain that colonizes to high density and causes acute otitis media, but typically does not cause invasive disease^38^. After three days of colonization, bacteria were recovered from the nasal passages via retro-tracheal lavage, allowed to expand overnight for 16 h on TSA-blood agar plates to recover bacterial populations, and subsequently harvested for genomic DNA extraction and re-infection of the next passage in the respective lineages. This methodology was repeated for a total of ten generations per mouse lineage, involving 30 total infections. Whole genome sequencing of the bacterial populations (to >300× coverage, enabling detection of any mutation at ~1% or greater) was undertaken at each passage and mutation frequencies were subsequently determined (Supplementary Table 1).

Following filtering of low-frequency mutations deemed unreliable or likely artifacts (see Methods), a final set of 782 mutations from 33 different samples was identified. The distribution of these mutations revealed several key features of this model of *S. pneumoniae* experimental evolution *in vivo*. First, the starting populations of the 19F strain used to initiate each of the three infections contained surprising genetic diversity: 97 instances of 68 unique mutations. These founding mutations ranged from 2-42% frequency and included many of the mutations that would ultimately be selected in subsequent transfers (Figure S1, Supplementary Data). Second, only ~16% of mutations detected during any given mouse infection were detected in any subsequent mouse (Figure S1). This demonstrates strong bottlenecks during the transfer protocol and establishment of the next infection, which involved both the subsampling of one-third of the population and a brief growth period on selective agar prior to inoculation. Third, once a given mutation persists to the next passage owing to a combination of selection and chance, its chance of being represented in a third passage is ~75% (Supplementary Data). At this point, these mutations encounter competition with other selected mutations and the presumably fitter genotype excludes the others. Ultimately, only 18 unique mutations were detected in six or more passages. To summarize, experimental evolution of *S. pneumoniae* during nasopharyngeal infections of the mouse exerts strong selection on a large pool of genetic variation, most of which is removed with each passage. This variation is replenished by new mutations (<10 / lineage) at each passage, but a few mutations undergo selective sweeps that exclude most of this variation. These sweeping mutations define the most likely adaptations in this environment.

To infer the genealogical structure of each population and visualize changes in lineage frequencies, we used the lolipop software package we developed that identifies linked genotypes and ancestry from shared mutation trajectories over time. These results are then displayed as Muller diagrams that integrate both frequency and ancestry (Figures 1A-C) and also as lineage-frequency plots (Figures 1D-F) or genetic pedigrees (Figure S2). Each of the three evolved populations were dominated by two genotypes, represented by green and orange in the plots, in which adaptive but ultimately unsuccessful genotypes (green) were outcompeted and replaced by a more fit lineage (orange) that nearly fixed in each population. These green and orange lineages subsequently diversified genetically and acquired new putatively adaptive mutations, which are represented by different colors nested within these green and orange backgrounds. We subsequently analyzed the identities and putative functions of these mutations below.

**Figure 1.**
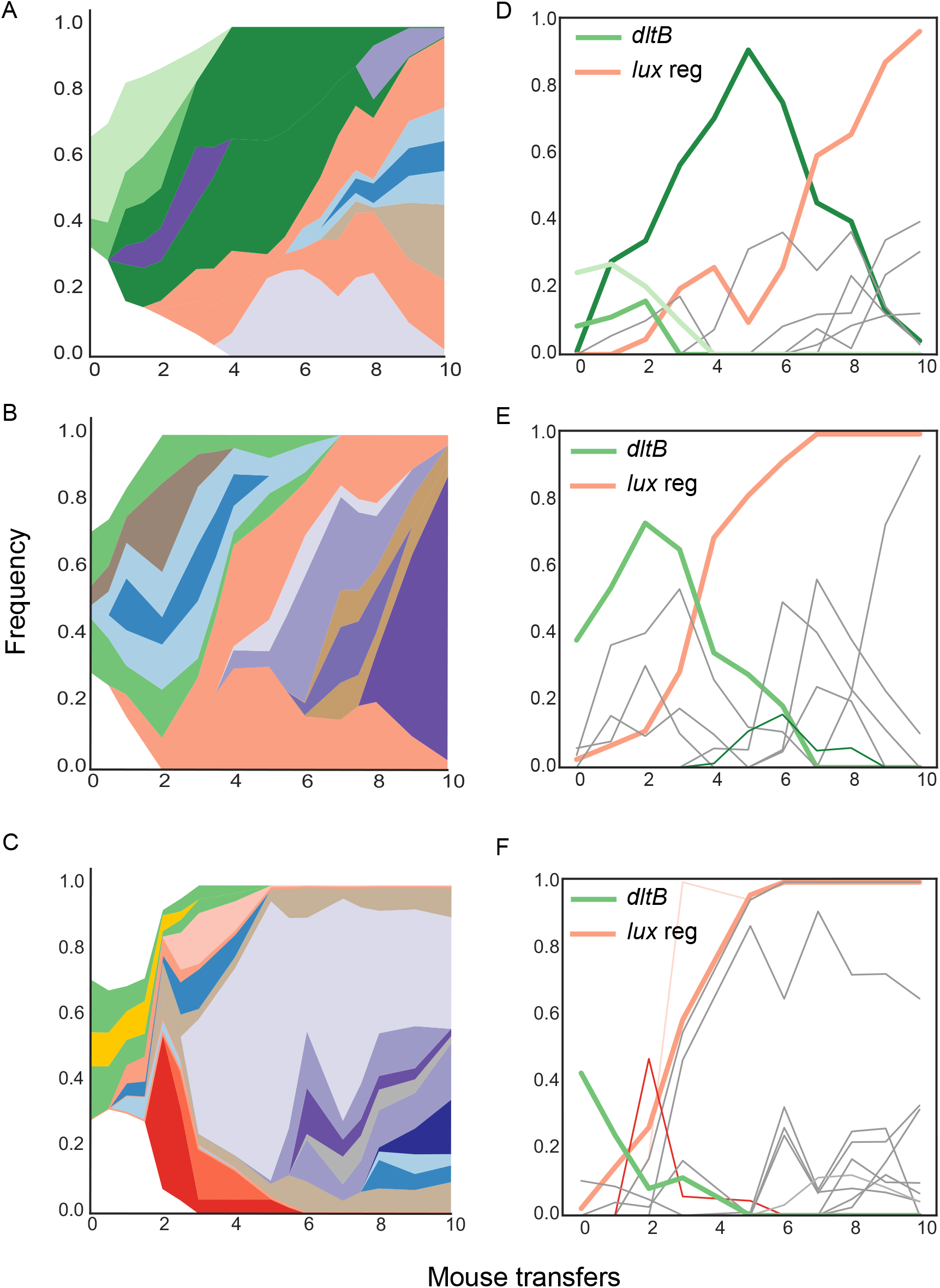
Evolutionary dynamics during repeated pneumococcal colonization. Each row corresponds to an independent evolution experiment (M1-M3). Muller plots of genotype ancestry and frequency (A-C) and individual genotype frequency plots (D-F) corresponding to each lineage. The two sets of primary driver mutations in *dltB* and the Lux transcriptional regulator are highlighted in green and orange, respectively. The red segments in panels C/F correspond to a *mutS*/*hexA* mutant genotype. Additional colors are arbitrary.

### Genes under selective pressure

After filtering for nonsynonymous and promoter mutations found at multiple consecutive time points, 56 mutations in 44 different genes were identified and are predicted to be have been under positive selection during experimental evolution in the mouse nasopharynx. One locus exhibiting the strongest evidence of positive selection, in which nonsynonymous mutations evolved in each lineage and reached >95% frequency, was a predicted helix-turn-helix (HTH) transcriptional regulator (Figure 2, Supplemental Table 1). The same mutation (L834V) swept in 2 of the populations because it was present in the shared inoculum, but its dynamics differed because of chance effects of other competing or linked mutations in these populations (Figure 2). The M1 population evolved a different mutation during the second infection cycle that ultimately reached high frequency by the end of the experiment (Figure 2). Further examination and targeted deletion of this mutant identified this gene as the main positive transcriptional regulator of the luminescent reporter locus that was engineered to enable Xenogen imaging, as both the evolved lineages and the targeted deletion of the gene resulted in a loss of luminescence (Lux). It seems likely that this loss of luminescence was selected to improve growth rate within the environment of the nasal passages, as the luminescence reaction was burdensome and drained cellular energy in the form of ATP. This was reflected by *in vitro* competitive index experiments, whereby the parental BHN97 strain outcompeted the bioluminescent BHN97x within two passages as measured by competitive index (Supplementary Figure 1). Targeted deletion of the HTH transcriptional regulator resulted in a fitness benefit in the BHN97x background, further establishing the mutation resulting in loss of bioluminescence was energetically favorable (Supplementary Figure 1). More significant, the fact that different mutations in this Lux regulator fixed in replicate populations indicates that their dynamics were driven by selection on new mutations rather than by genetic drift, recombination, or high mutation pressure. Further, their varied trajectories demonstrate effects of these other population-genetic processes, including contributions of other selected mutations on the HTH background and on competing lineages (Figure 2). As the loss of the marker is irrelevant to specific fitness in the nasopharynx, subsequent investigations focused on additional loci that appeared to be under strong positive selection.

**Figure 2.**
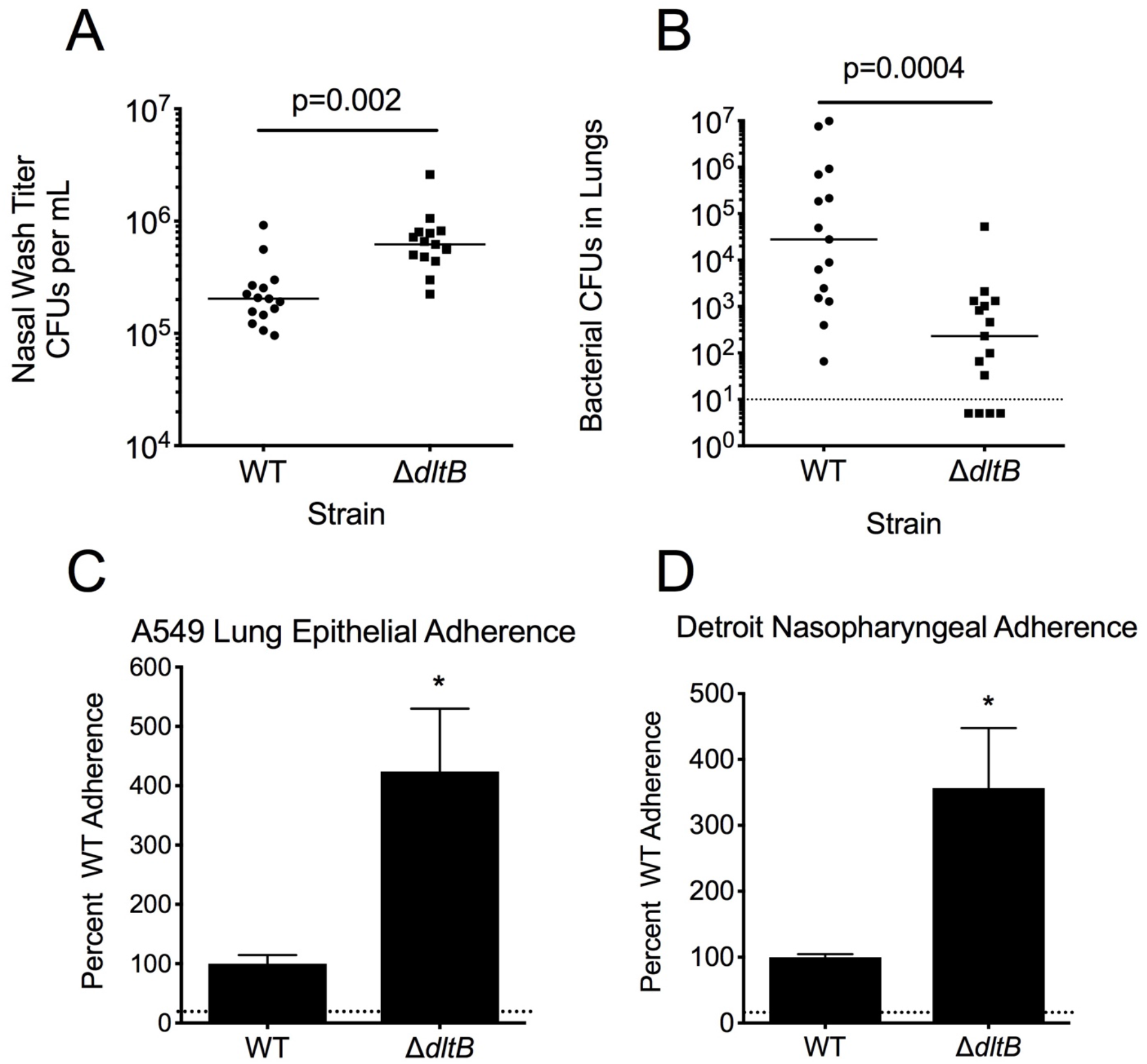
Impact of loss of function of *dltB* on respiratory infection and host cell interaction. Bacterial titers of the parental and *dltB* insertional mutant 72 h post-challenge in the nasal lavage (A) and the lung homogenates (B). Each data point represents an individual mouse. Relative adherence of the parental and *dltB* insertional mutant to A549 lung epithelial (A) and Detroit nasopharyngeal (B) cell lines. Adherence assays were performed in biological triplicate. * indicates p<0.05 by Mann-Whitney.

A frameshift mutation in *hexA* (the *mutS* ortholog) was observed, generating a hyper-mutator subpopulation that transiently dominated the M3 population for one generation, but was subsequently lost in later passages (Figure 1C, red lineage). Deletion of this gene did not result in any discernible fitness benefit during infection in either the lungs or the nasal passages (Supplemental Figure 2A), but greatly increased the mutational frequency of the pneumococcus (Supplemental Figure 2B). These data indicate that hyper-mutator strains have the potential to emerge *in vivo*. However, other better adapted lineages may outcompete them if beneficial mutations are not acquired.

The gene *dltB*, which showed strong evidence of selection, then became the focus of our investigation. This locus is involved in the D-alanylation of the teichoic acids of Gram-positive bacteria. DltB is proposed to be involved in surface charge modulation and contributes to resistance against cationic antimicrobial peptides and antibiotics in multiple species of Gram-positive bacteria^39–42^. The starting populations contained seven distinct frameshift mutations in the *dltB* locus, with four new mutations arising in later time points in multiple lineages (Figure 2, Supplemental Table 1). The acquired mutations were predicted to abrogate protein function and so we conducted investigated an isogenic *dltB* deletion mutant to discern the factors driving emergence of these mutants during repeated nasal colonization.

### Role of *dltB* in Host Cell Adherence

Colonization of host tissues by bacteria is mediated by the binding of adhesion molecules to surface molecules on resident host cells. However, this process is typically first initiated by physicochemical forces that include Van der Waals interactions, acid–base hydrophobic interactions, and electrostatic charge^43^. In the pneumococcus, surface charge is predominantly dictated by the capsule. However, modification of cell wall teichoic acids is another mechanism by which the bacterial surface charge can be modulated. This has previously been shown in *Entereococcus faecalis*, where adherence to host cell surfaces was enhanced by mutation of *dltA*^44^. Here, we hypothesized that mutation of pneumococcal *dltB* could be advantageous by preventing teichoic acid D-alanylation. This would alter the pneumococcal surface charge and increase host cell adherence. Here, we addressed this hypothesis using human cell culture models of both nasopharyngeal and lung epithelial cells. Adherence to both Detroit nasopharyngeal and A549 lung cells was significantly enhanced in the *dltB* mutant strain by comparison with the parental strain (Figure 3, A-B). Taken together, these data indicate that loss of D-alanylation of the teichoic acid provides a selective advantage for the initial adherence of *S. pneumoniae* to host cells.

**Figure 3.**
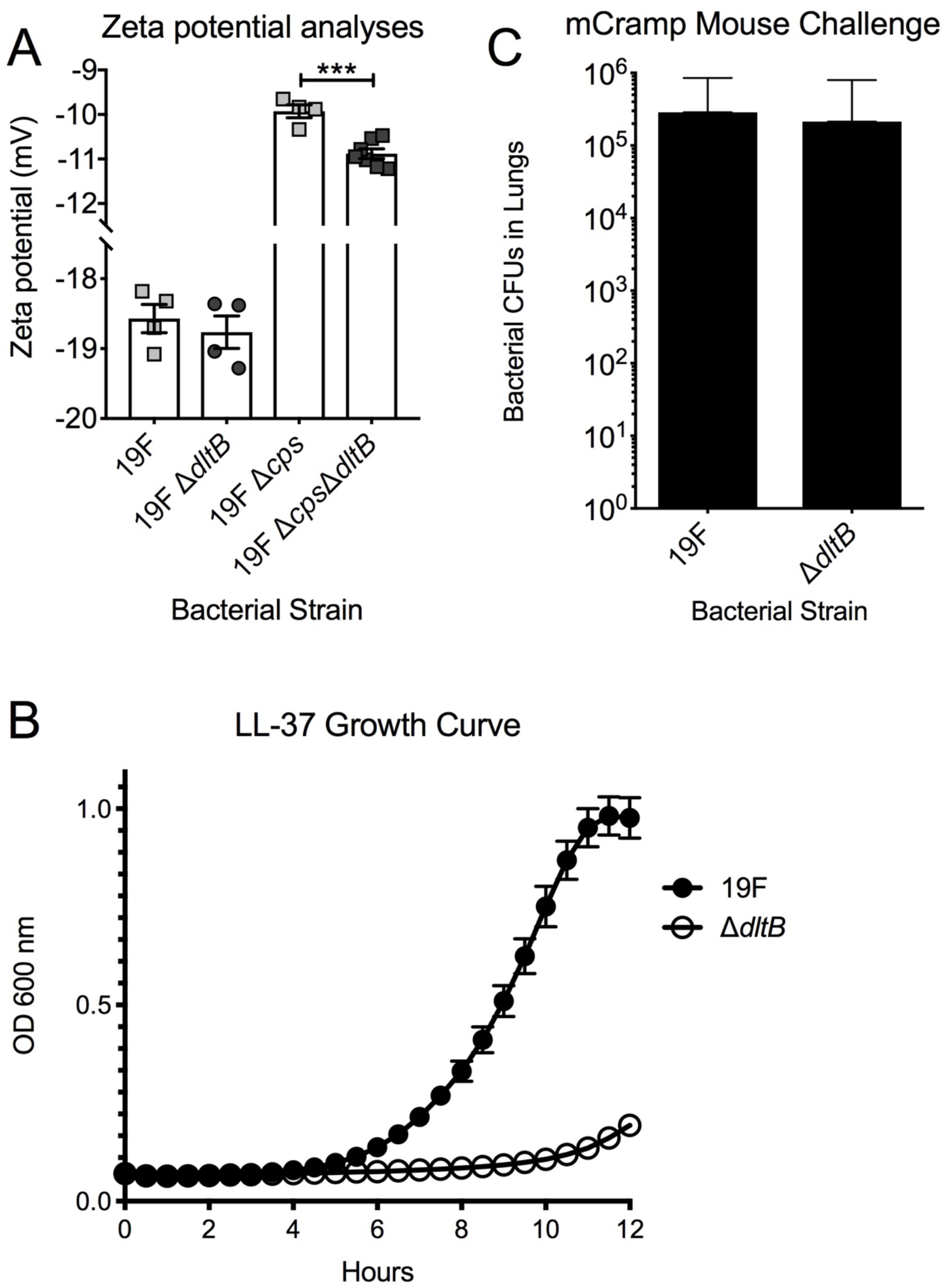
Impact of loss of function of *dltB* on surface charge and sensitivity to antimicrobial peptides. Measurement of zeta potential of the parental wild-type and *dltB* mutant strain both with and without capsule expression (A). Growth inhibition of Δ*dltB* by supplementation of media with sub-inhibitory concentrations of LL-37 (B). Bacterial CFUs recovered from murine lungs of mCramp mice demonstrated no discernible difference between the parental BHN97x and the Δ*dltB* following challenge (C).

### Role of *dltB* in Murine Nasal Colonization

Building on the above findings, we hypothesized that the *dltB* mutant strain would have a competitive advantage in murine colonization due to its enhanced adherence relative to the wild-type. We observed that deletion of *dltB* provided a significant advantage in the murine nasal lavage, with bacterial burden significantly greater at 72 h post-challenge (Figure 3A). In the same challenge experiment, the relative bacterial burden in the lungs was measured. In contrast to the nasal titer results, deletion of *dltB* conferred a significant disadvantage in terms of bacterial burden in the lungs at three days post intratracheal challenge (Figure 3B). These data suggest the loss of *dltB* is favored by selection during nasal colonization, but this fitness benefit was niche specific with a corresponding fitness tradeoff when bacteria were administered directly into the lungs.

### Role of D-alanyation on Surface Charge

The main determinant of *S. pneumoniae* surface charge is the polysaccharide capsule. Capsules of greater negative charge have been shown to impart a selective advantage during nasal colonization^26^. However, modification of the teichoic acids by D-alanylation alters the bacterial surface charge^45^. Accordingly, we sought to determine if deletion of *dltB* conferred an impact on the net surface charge by zeta potential measurement. No discernible differences were observed in the encapsulated bacteria, with the zeta-potentials reflective of serotype 19F strains from previous reports (Figure 4A) ^46^. While capsule is a requirement for invasive disease, previous studies have indicated capsule can be actively shed in response to inflammation and that non-encapsulated mutants can colonize the mucosal surface, albeit with significantly reduced densities ^37,47^. Accordingly, we examined the impact of the *dltB* deletion on surface charge in both the presence and absence of the polysaccharide capsule. Using mutant strains deficient in both *dltB* and the capsule locus we observed that non-encapsulated bacteria had a significant net increase in the negative charge of the bacterial surface (Figure 4A). Therefore, we propose that evolved defects in *dltB* modulate teichoic acids and results in altered surface charge properties consistent with enhanced bacterial adherence.

**Figure 4.**
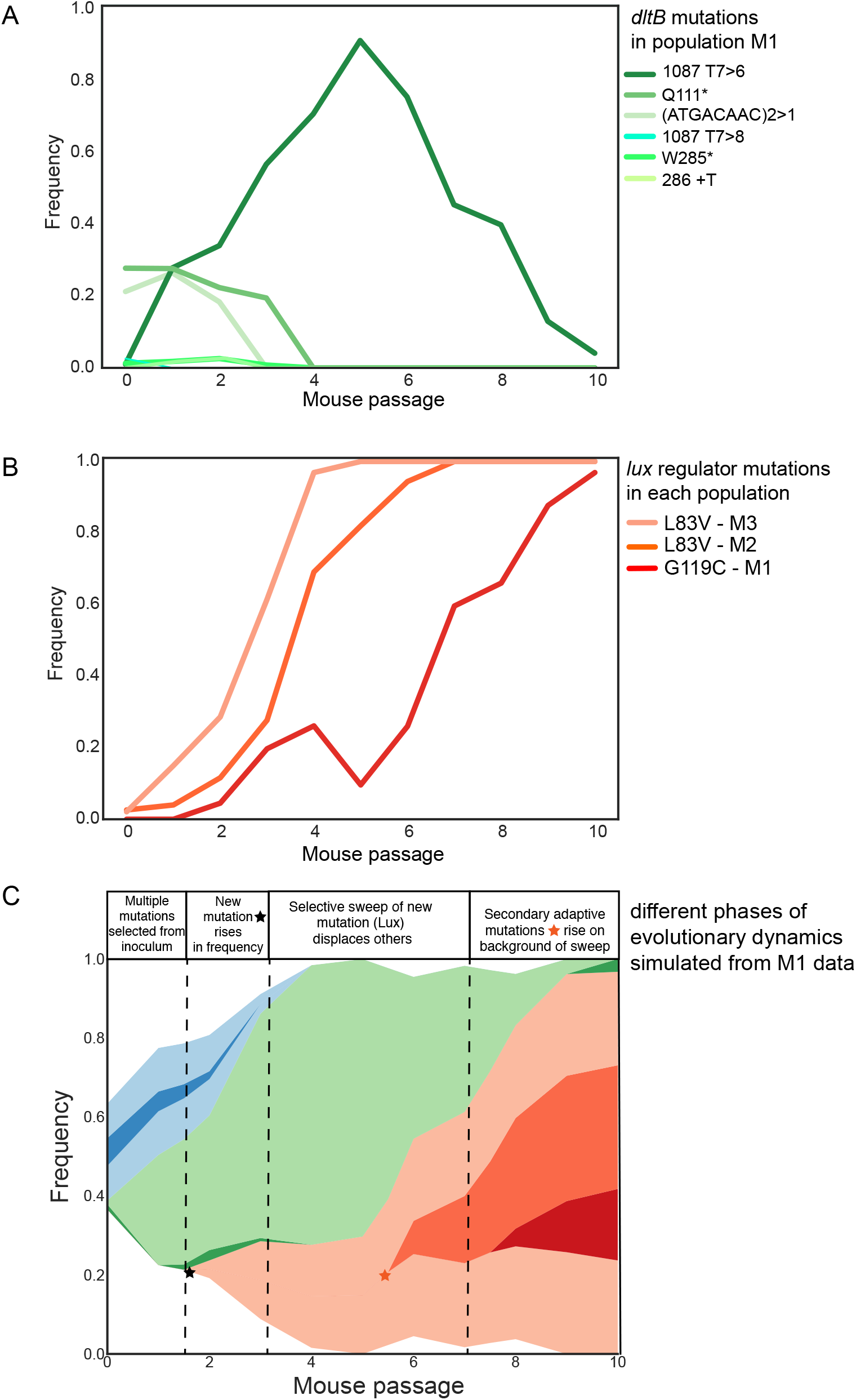
Summarized population-genetic dynamics of selected mutations. A. Dynamics of contending *dltB* mutations found in population M1. B. Dynamics of mutations eliminating luciferase production in each population. C. Muller plot of the interaction between these selected genotypes and the eventual rise of secondary adaptive mutations on the mutated Lux genetic background.

### Role of *dltB* in Antimicrobial Peptide Resistance

The fitness benefit of the *dltB* mutant in the nasal passages was balanced by a fitness defect in the lungs. One potential explanation for this difference is that the innate immune response to pneumococcal lung infection is robust, with significant cellular infiltrate and corresponding increase in the concentration of cationic antimicrobial peptides during infection^48^. D-alanylation of teichoic acids has previously been shown to be important for the capacity of bacteria species to resist the bactericidal activity of positively charged antimicrobial peptides^45^. In addition to the bactericidal activity, antimicrobial peptides can also induce enzymatic release of the polysaccharide capsule from the pneumococcal surface^37^. To distinguish between these two possibilities, we examined whether deletion of *dltB* impacted the capacity of the pneumococcus to release capsule in a LL-37 dependent manner. In response to sublethal LL-37 exposure, no significant difference in polysaccharide capsule shedding was observed as determined by capsule blotting with 19F specific antisera (Supplementary Figure 2).

As there were no discernible differences observed in capsule shedding, we next sought to determine if the deletion of *dltB* resulted in differential sensitivity to the antimicrobial peptide LL-37. Supplementation of media with varying concentrations of LL-37 revealed that *dltB* mutant had heightened sensitivity to the antimicrobial peptide (Figure 4B). We hypothesized that this sensitivity was one of the primary mechanisms underlying the fitness defect of the *dltB* mutant in the murine lungs. To address this, we undertook murine challenges in mCRAMP, a cathelicidin-related antimicrobial peptide deficient animal. This absence of a major cationic antimicrobial peptide in this murine background results in failure to induce pneumococcal capsule shedding and increased susceptibility to bacterial challenge^37^. Here, we observed that the bacterial burden in murine lungs was not significantly different between the parental and the *dltB* mutant strain (Figure 4C). Hence, these data show that the selective benefits arising from loss of *dltB*, that is increased adherence and nasal colonization, are counterbalanced by an increased susceptibility to antimicrobial peptides during acute inflammation.

### Additional traits under selection *in vivo*

Mutations in other genes experienced positive selection during this experiment. Despite a high level of genetic variation (25-40 detected mutations) in the founding population, ~60 new mutations arose across the three replicate lineages that were undetected in the inoculum. This indicates strong selection on a variety of traits during the *in vivo* evolution experiment (Figure S2, Supplementary Data). Some of these mutations may have been enriched by linkage to another mutation under selection, producing the genotypes depicted in Figure 1. However, others suggest at selected traits beyond D-alanylation. For example, mutations that fixed include nonsynonymous mutations in *briC*, encoding biofilm-regulating peptide, in *tig*, encoding trigger factor, and in a gene encoding an ABC transporter putatively involved in thiamin biosynthesis (orthologous to TIGR4 SP2197). All three mutations fixed because of their linkage to the HTH mutation that eliminated Lux production, so they could be simply neutral or mildly deleterious ‘hitchhiking’ mutations. Nevertheless, it is also possible they contributed to genotype fitness. Mutations that rose to high frequency independently of this Lux regulator included a mutation in *copA*, encoding a copper exporter, in a promoter directly upstream of the PsaABC manganese importer, in *pdhC* encoding a component of pyruvate dehydrogenase, and in a gene encoding an arylsulfatase enzyme. Interestingly, the *copA* and arylsulfatase mutations comprised a linked genotype that appears to have avoided being excluded by the selected Lux genotype by a recombination event that combined these mutations (Figure 1F). Likewise, *pdhC* and *psa* promoter mutations comprise a lineage that nearly fixed following the sweep of the Lux genotype (Figure 1E). These genotypes therefore provide the strongest evidence of selection on traits independent of *dltB* or Lux and indicate that the transport of metals and central metabolism experience selection during serial passage in the murine nasopharynx.

## Discussion

We developed an experimental model of *in vivo* evolution of *S. pneumoniae* with two major objectives: first, to identify adaptive genotypes and traits during colonization of the mouse nasopharynx; and second, to discover the population-genetic dynamics of repeated pneumococcal colonization. At the outset, we considered whether the experimental cycle of repeated mouse infections might produce severe population bottlenecks that would limit the power of selection and inflate the role of genetic drift. Strong selection is predicted to produce evolutionary parallelism, with mutations in similar genes and the same traits evolving in the same conditions. In contrast, strong bottlenecks can oppose selection and diminish repeatability by increasing effects of chance. Yet despite only three replicate lineages, we observed substantial gene-level parallelism in selected mutations causing similar or identical trait changes.

Although the shared selective sweep eliminated the luminescence marker that had been introduced for *in vivo* imaging, only serving to demonstrate its fitness cost, other mutations reaching high frequency in advance of or following this sweep show that selection on putatively host-adaptive traits can outweigh effects of mutation and drift in this model. Further, despite the loss of genetic diversity during each manual transmission between mice, many mutations persisted through the infection cycle or arose *de novo* within each infection to fuel subsequent evolution. This study, reinforces evidence from recent publications, that demonstrates that the mutated genes are important for pneumococcal colonization and persistence within the nasopharynx.

Most significantly, we identified parallel loss-of-function mutations in *dltB* that increased pneumococcal cellular adherence and generated a pronounced fitness advantage during colonization (Figure 3). It is noteworthy that several of these *dltB* mutations were frameshifts caused by expansions or contractions in sequence repeats, and most were present at low frequency in the initial inoculum, which implies that this locus may have evolved to readily produce potentially adaptive, heritable variation as a contingency locus ^49^. These results indicate that polymorphism in *dltB*-associated phenotypes may be present in many pneumococcal populations and influence colonization dynamics. Further, had mutations eliminating luciferase production not produced even greater fitness advantages, *dltB* mutations would have likely fixed in all populations because of their selective benefit.

While the polysaccharide capsule locus is a very genetically and biochemically diverse virulence factor, other genetic loci like *dltB* may be subject to diversifying selection as well. Clinical isolates of *S. pneumoniae* are highly variable in their sensitivity to antimicrobial peptides, with both capsule type and genetic background playing important roles in the relative sensitivity of strains to this host defense mechanism^50^. Colonization and transmission of *S. pneumoniae* are critical mediators of the success of the pneumococcus and represent strong evolutionary pressures on this human pathogen. The evolution experiment indicated that loss of D-alanylation of the teichoic acids conferred a fitness advantage during interactions with host epithelial cells and during nasal colonization.

As colonization is a major aspect of pneumococcal fitness, the selective conservation of this gene in *S. pneumoniae* was cast into question. We hypothesize this gene is maintained because its deletion is associated with increased sensitivity to antimicrobial peptides and thus reduces fitness in niches with significant inflammatory infiltrate, such as the pneumonic lung. In addition, recent studies have also demonstrated a role of *dltB* in pneumococcal transmission whereby mutants in this locus have significantly reduced shedding from colonized mice which reduces mammalian transmission^10^. The increased adhesion and colonization phenotypes conferred by loss of this locus may initially be beneficial, but the failure to transmit to new hosts may explain the retention of this gene in *S. pneumoniae* given the fitness advantages we observe during colonization. The serotype 19F capsule type, which was the strain utilized in these studies, is one of the most negatively charged capsule types and this electronegativity is thought to confer a fitness benefit during co-colonization with multiple serotypes^46^. While the change in surface charge was masked by the polysaccharide capsule, recent studies have indicated that pneumococci can enzymatically release their surface polysaccharide in response to antimicrobial peptides both *in vitro* and *in vivo^37^*.

The experiment also revealed several other mutations of potential functional significance to *S. pneumoniae* colonization (Figures 1, 4, S2). These include an altered allele of the small peptide BriC that regulates biofilm production, recombination competence, and colonization^51^, and four mutants affecting metabolism, including thiamin uptake, copper efflux, manganese uptake, and pyruvate dehydrogenase. These findings add to considerable evidence that the pneumococcus experiences selection for efficient trace metal uptake and glycolytic flux during host colonization^52^. The specific contributions of these mutations to fitness in the nasopharynx remain to be determined but their rapid rise in frequency independently of *dltB* and Lux regulator mutations almost certainly reflect strong selective advantages^53^.

This study demonstrates that serial infections that maintain experimental bacterial populations can generate strong selection for mutations that enhance colonization or pathogenicity. This is by itself not surprising; however, the most common analysis method would involve WGS of randomly selected clones at the conclusion of the experiment, which would likely only reveal the Lux regulator as the selected mutational target. This result alone might be judged a failure. However, our use of whole-population genomic sequencing to a depth sufficient to capture mutations >1% frequency or greater revealed many more genetic targets as well as the population dynamics of the infection cycle. These additional targets demonstrate a surprising degree of predictability in the bacterial population dynamics in this infection cycle that preserved genetic variation over time. We were concerned *a priori* that the bottleneck imposed during the transfer process from mouse to mouse would diminish the effectiveness of selection in the experiment, with each new host producing strong founder effects and the loss of genetic variation. This was not the case, as each new population sample clearly resembled the previous one of that lineage with only modest sampling artifacts. The genetic parallelism in *dltB* and lux mutations also illustrate the predictability and reproducibility of the study system.

We suggest that experimental evolution can be readily applied to multiple *in vivo* model systems to understand host adaptation with an eye toward specific host niches^54,55^. This approach demonstrates that alterations of teichoic acid can enhance pneumococcal fitness by improving nasal colonization and host cell adherence. However, these adaptive mutations were highly niche-specific and imposed a tradeoff in the ability to withstand host defenses, reducing fitness during deeper tissue invasions such as lung infection and increasing sensitivity to antimicrobial peptides. These results suggest the *dltB* gene is subject to balancing selection to meet these different functions and may explain why it has evolved a higher mutation rate caused by repetitive sequences, so that most populations harbors a *dltB* variant suitable to the given niche. These methods hold promise for the study of other host-microbe interactions to identify tissue tropisms or colonization strategies.

## Acknowledgements

JWR and VC are supported by 1U01AI124302. JWR is supported by 1R01AI110618. This work was supported by St. Jude Children’s Hospital and ALSAC. This work was supported by the Australian Research Council (ARC) Discovery Project Grant DP170102102 to JWR and CAM. SLN is a National Health and Medical Research Council Early Career Research Fellow (1142695) and CAM is an ARC Future Fellow (FT170100006).

## Supplementary Figure Legends

**Figure S1.**
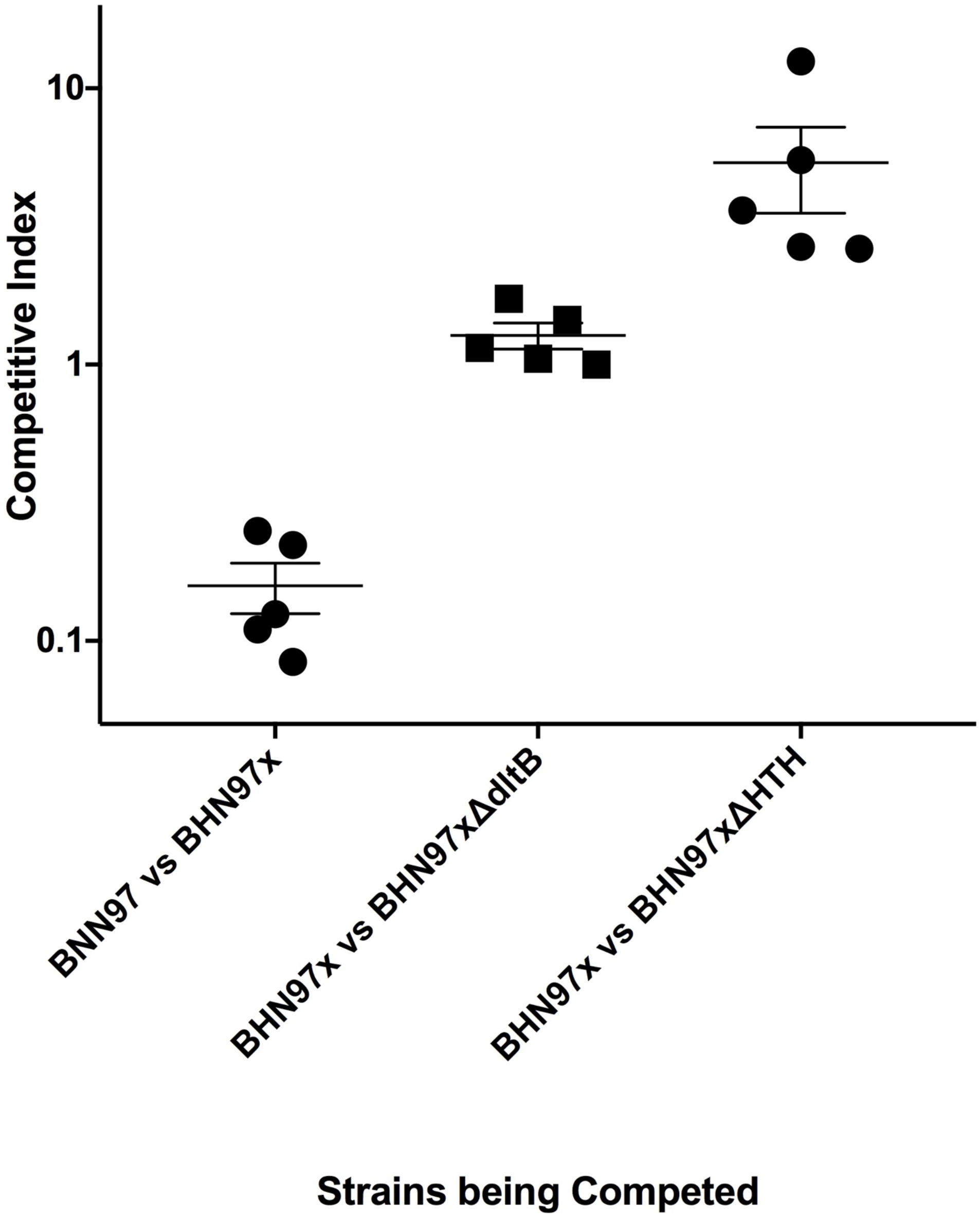
Fitness benefit for loss of bioluminescence expression. Strains were directly competed against either other under *in vitro* conditions for two passages of approximately 12 generations each. Data values are for the relative fitness of the second strain versus the first on the x-axis. Introduction of the lux cassette (BHN97x) resulted in a fitness defect when competed against the parental BHN97. A loss of function mutation in the helix turn helix regulator whose deletion results in loss of expression of the lux cassette likewise results in a fitness benefit against the respective parental BHN97x. No discernible benefit for the loss of *dltB* was observed during *in vitro* growth. Each data point indicates and independent lineage.

**Figure S2.**
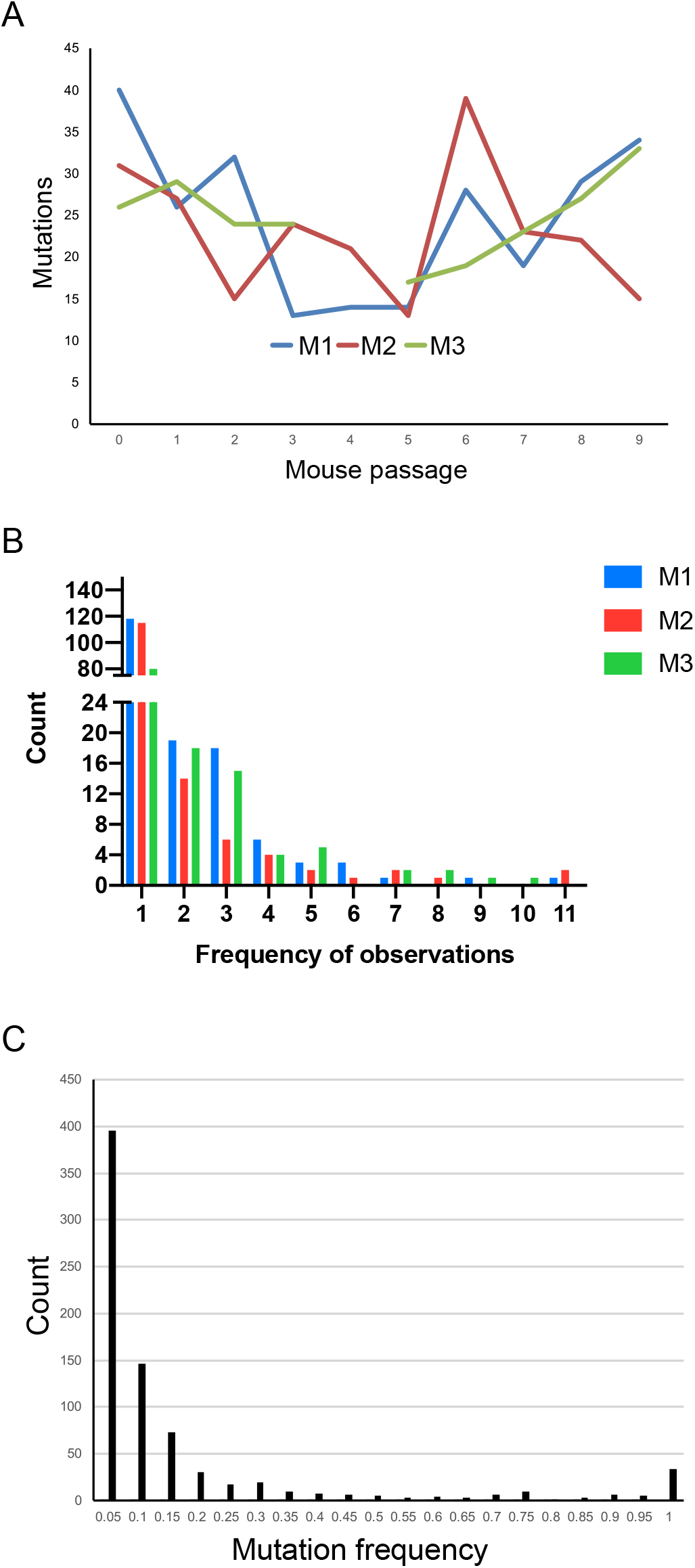
A. Number of mutations per lineage identified in the inoculum (0) and following each mouse passage, following filtering. The sample for M3, passage 4 did not pass quality filtering. **B.** Histogram of the number of observations of mutations detected in each lineage (M1-M3). Most mutations were detected only once but many (12-18) were detected in two or more samples. **C**. Histogram of all mutations by their observed frequency in a given sample, including repeated observations of the same mutation.

**Figure S3.**
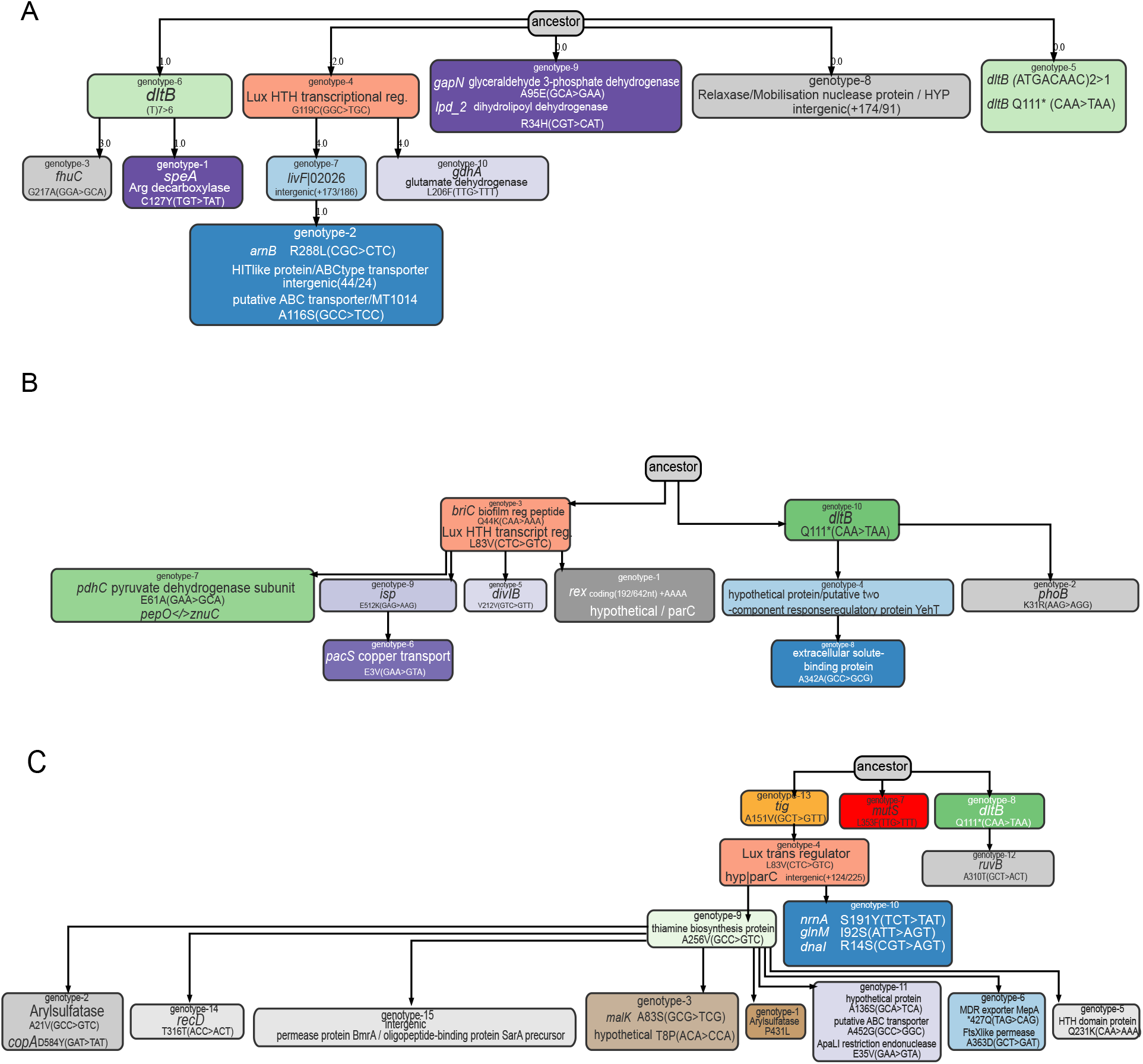
Genotype ancestry diagrams for each evolved lineage, as inferred by the lolipop software package. Each box indicates new mutations inferred in history of that lineage. The presence of multiple mutations in some evolutionary steps results from uncertain order of evolution based on population-sequencing data.

Table S1. Mutations identified in each population that were filtered for those likely under selection and used to infer lineage ancestry and frequency. Identified mutations, their relative frequency, and the lineage/passage they were detected are detailed herein. A second tab summarizes mutations that comprise the trajectories shown in Figure 1.

